# Continuous vibration-driven virtual tactile motion perception across fingertips

**DOI:** 10.1101/2022.09.06.506303

**Authors:** Mehdi Adibi

## Abstract

Motion perception is a fundamental function of the tactile system, essential for object exploration and manipulation. While human studies have largely focused on discrete or pulsed stimuli with staggered onsets, many natural tactile signals are continuous and rhythmically patterned. Here, we investigate whether phase differences between *simultaneously* presented, *continuous* amplitude-modulated vibrations can induce the perception of motion across fingertips. Participants reliably perceived motion direction at modulation frequencies up to 1Hz, with discrimination performance systematically dependent on the phase lag between vibrations. Critically, trial-level confidence reports revealed the lowest certainty for anti-phase (180°) conditions, consistent with stimulus ambiguity as predicted by the mathematical framework. I propose two candidate computational mechanisms for tactile motion processing. The first is a conventional cross-correlation computation over the envelopes; The second is a probabilistic model based on the uncertain detection of temporal reference points (e.g., envelope peaks) within threshold-defined windows. This model, despite having only a single parameter (uncertainty width determined by an amplitude discrimination threshold), accounts for both the nonlinear shape and asymmetries of observed psychometric functions. These results demonstrate that the human tactile system can extract directional information from distributed phase-coded signals in the absence of spatial displacement, revealing a motion perception mechanism that parallels arthropod systems yet potentially arises from distinct perceptual constraints.

## Introduction

Motion is a fundamental quality of sensory input. In vision, despite diverse evolutionary trajectories across different species, from insects and cephalopods to vertebrates, visual systems have converged on fundamentally similar mechanisms of motion processing [1, 2]. However, motion is not exclusive to vision; it is also a hallmark the tactile sensory system, with considerable behavioural relevance for both animals and humans.Everyday interactions such as object manipulation and haptic exploration involve relative motion between the skin and surfaces [3]. For example, discerning roughness and smoothness, identifying material properties (e.g. metal vs. wood) or recognising object shapes requires dynamic contact through palpation and movement. Reading Braille depends on lateral movement of the fingertips to interpret sequences of raised dots. Tactile motion processing underpins fine motor control and precise object manipulation [4, 5]. Yet, the perceptual and computational mechanisms underlying tactile motion remain poorly understood.

Two principal sources of information for tactile motion have been identified in the literature [5, 6]. The first relies on the sequential activation of mechanoreceptors at different skin locations as an object or stimulus moves across the skin – such as when an insect crawls along the arm. This includes apparent tactile motion, typically studied experimentally using discrete, pulsed stimuli delivered in succession to separate skin sites [7–14]. The second source involves skin deformation cues, particularly shear and stretch, which arise during sliding contact or friction. These deformations can convey directional information [15], potentially through recruitment of distinct afferent populations, such as slowly adapting type II (SA2) units, which are sensitive to stretch and contribute to motion direction perception [16].

At the level of periphery, slowly adapting type I (SA1) afferents convey high-resolution spatial information about contact location [4, 17, 18]. The spatiotemporal pattern of their population activity is thought to encode motion direction and speed with acuity comparable to human perceptual performance [18]. Rapidly adapting type I (RA1) afferents may also contribute to motion encoding via spatiotemporal patterns, though with lower spatial precision due to their larger receptive fields [18]. SA2 afferents, previously mentioned, respond to skin stretch and support motion direction perception through their tuning to deformation patterns [15, 16].

Here, I investigate a third and less explored mechanism for tactile motion: the perception of motion across fingers induced by asynchronous, continuous streams of vibrotactile input delivered to two fingertips simultaneously. Unlike prior studies that typically rely on either (a) sequential, pulsed stimulations delivered to multiple discrete skin sites [7, 9, 11, 14, 19], or (b) physical sliding stimuli that engage friction-induced skin deformation [5, 6], this paradigm involves two spatially fixed but temporally dynamic inputs. The stimulation does not involve skin movement or high spatial acuity, but rather evokes motion percepts through temporal phase differences between inputs to two fingerpads.

Fingertips are among the most densely innervated tactile regions in the mammalian and human body [20], and are primary organs for active exploration [3]. This work uses continuous amplitude-modulated vibrations known to predominantly recruit rapidly adapting type II (RA2) afferents – i.e., Pacinian corpuscles – which are sensitive to high-frequency vibratory energy and exhibit large receptive fields [21–23]. Unlike SA1 and RA1 afferents, RA2 units are less sensitive to fine spatial features but can detect remote vibratory events across skin and even bone. Notably, the stimuli here are delivered over the entire fingertip pad, eliminating reliance on fine spatial localisation and instead leveraging temporal synchrony or asynchrony across digits. This approach is analogous in principle to mechanisms found in certain arthropods, such as chelicerates, which detect and localise remote vibratory sources using their paired appendages [24]. Similarly, humans may infer motion direction or location of a remote source by comparing asynchronous vibratory input across fingerpads [25] – effectively extending tactile spatial perception beyond the point of contact.

In this study, I first formalise the physical basis for detecting the location and direction of a remotely moving vibration source, and how such stimuli can be simulated through asynchronous amplitude-modulated input to two fingertips. I then present a series of psychophysical experiments characterising human perceptual performance in detecting the direction of such inferred motion, revealing a novel mode of tactile motion perception that operates independently of spatial acuity or physical movement between surface and skin.

## Materials and methods

Vibrotactile stimulation is a versatile method for conveying spatiotemporal information through the skin and has been widely employed in both fundamental research and haptic technology applications [26]. Arrays of actuators delivering temporally staggered pulses have been used to generate apparent motion across body surfaces such as between the hands, along the arm, or over the back, simulating a moving tactile stimulus without physical displacement [8, 12, 19, 26, 27]. While such paradigms rely on discrete bursts or pulses with differences in stimulus onset timing across spatially distinct sites to evoke motion percepts, the current study employs continuous amplitude-modulated waveforms with controlled phase offsets, enabling investigation of motion perception from contentious, distributed, phase-coded signals in the absence of distinct onsets or spatial displacement.

To model the stimuli generated by a remotely vibrating source, such as a mobile phone on a table, we consider a point source emitting a sinusoidal carrier wave *A*_0_ sin (2*πf*_*c*_*t*), where *A*_0_ and *f*_*c*_ denote the amplitude and carrier frequency of vibration, respectively. As this vibration propagates through the substrate – e.g., the table – approximately as a plane wave, it undergoes attenuation due to dissipation, scattering, and absorption. This attenuation typically follows an exponential decay with distance:

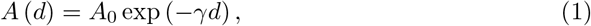

where *γ* is the attenuation coefficient (dependent on medium properties and frequency), and *d* is the distance from the source.

When the source moves relative to a fixed point, the amplitude at that point changes with time due to the variation in distance. These changes are proportional to the radial component of the source’s motion. In the special case of periodic movement at a frequency *f*≪*f*_*c*_, the received signal envelope at a fixed remote point with time-varying distance *d* (*t*) is itself periodic, and the received signal can be written as:

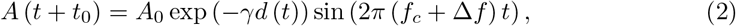

where *t*_0_ is the delay due to propagation over distance *d* (*t*), and Δ*f* is the Doppler shift caused by the source’s motion. Both parameters *t*_0_ and Δ*f* depend on the wave propagation speed in the medium, which is determined by its stiffness and density (e.g., ∼5790 ms^-1^ in stainless steel, and ∼3960 ms^-1^ in hard wood).

For distances on the order of a meter or less, *t*_0_ corresponds to sub-millisecond or nanosecond delays, well below biologically plausible detection thresholds. Moreover, assuming slow motion of source relative to wave propagation speed, and *f* ≪ *f*_*c*_, the Doppler shift Δ*f* is negligible. Henceforth, I assume *t*_0_ ≈ 0 and Δ*f* ≈ 0.

## Motion direction estimation via two touch points

When a sensor (e.g., a fingertip) is placed at a fixed point (hereafter a ‘touch point’), the direction of source movement along the radial axis (toward or away from the touch point) can be inferred from temporal changes in the vibration envelope. However, a single touch point provides no information about the tangential component of the motion. To recover trajectory information, at least two touch points positioned at distinct spatial locations are required (Fig. 1).

**Fig 1.**
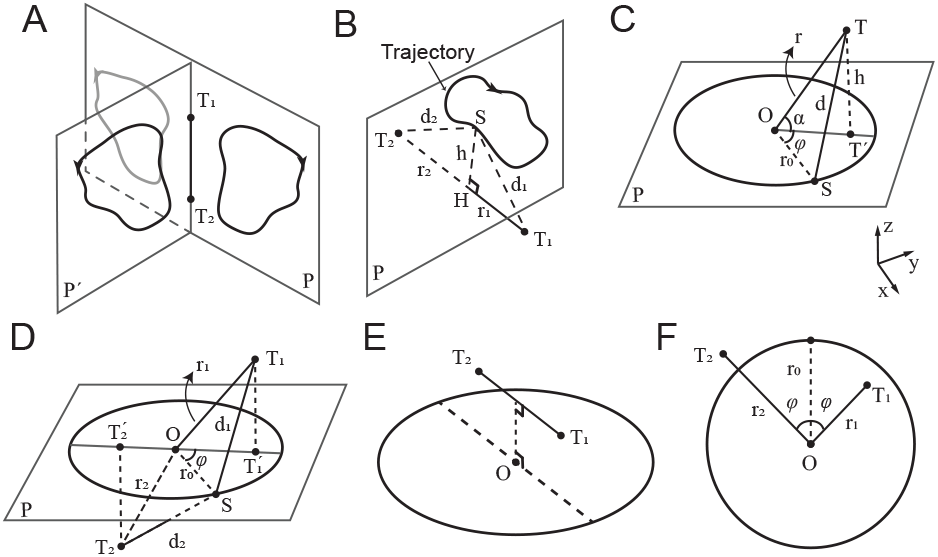
Detecting the motion of a remote vibrating source through patterns of vibrations sensed at two touch points. **A**, *T*_1_ and *T*_2_ denote the two touch points. The trajectory on plane *P*′ is the rotation of trajectory on plane *P′* around the touch axis *T*_1_*T*_2_. The grey closed curve shows the mirror of the trajectory with respect to the touch axis *T*_1_*T*_2_. **B**, For any arbitrary trajectory on the plane *P*, when the touch axis *T*_1_*T*_2_ is orthogonal to *P*, the vibrations from source *S* received at *T*_1_ and *T*_2_ are in-phase. *r*_1_ and *r*_2_ represent the distances of *T*_1_ and *T*_2_ from *P* respectively. *d*_1_ and *d*_2_ denote the distances from the source *S* to *T*_1_ and *T*_2_ respectively, and vary as *S* moves along the trajectory. **C**, A circular trajectory with radius *r*_0_, centred at *O. T* ′ denotes the projection of touch point *T* onto plane *P*. *h, r* and *d* denote the distances from *T* to *P, O* and *S*, respectively. *α* is the angle between *OT′* and *OT*′. **D**, *d*_1_ and *d*_2_ denote the distances from the source *S* to *T*_1_ and *T*_2_, respectively, and vary as *S* moves along the trajectory. *r*_1_ and *r*_2_ are the distances from *O* to *T*_1_ and *T*_2_, respectively. **E**, An example of anti-phase vibrations, when the projection of the axis *T*_1_*T*_2_ (dashed line) onto the trajectory plane *P* passes through *O*. **F**, The two-dimensional geometry. All conversions as in **D**.

Let *T*_1_ and *T*_2_ denote two such points. The envelopes of vibration received at these two locations can, in principle, be used to infer the moment-by-moment position of a moving source in 2D. However, the reconstruction is ambiguous: any trajectory and its mirror reflection across the line connecting the two touch points (the touch-point axis) yields identical vibration patterns. This is illustrated in Fig. 1A.

In 3D, ambiguity increases: all trajectories that are rotationally symmetric around the touch-point axis produce indistinguishable vibration profiles at the two points. That is, any trajectory that can be rotated about this axis into another remains perceptually equivalent at the touch points, leading to infinite number of trajectories that create an identical vibration pattern at the two touch points *T*_1_ and *T*_2_.

### In-phase vibrations

As discussed, a periodic source movement with frequency *f* results in a periodic amplitude modulation at each touch point. If the envelopes at *T*_1_ and *T*_2_ vary together over time – i.e., they are monotonic transformations of one another under a strictly increasing odd function – they are said to be ‘in phase’. For example, consider a trajectory confined to a plane perpendicular to the touch-point axis (Fig. 1B). The closest and farthest positions on the trajectory to *T*_1_ are identical to those to *T*_2_. Let *h* (*t*) denote the instantaneous orthogonal distance from the touch-point axis to the source trajectory at any moment *t*, and let *r*_*i*_ be the perpendicular distance from touch point *T*_*i*_ to the plane of the trajectory. The amplitude at each touch point is given by:

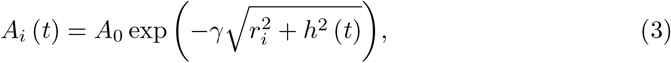

and the derivative:

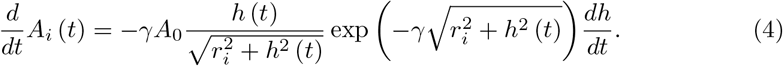

This shows that the envelope at each touch point changes in the same direction (increasing or decreasing together), confirming they are in phase. Additionally, one can show:

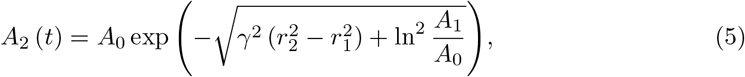

which is a strictly increasing function of *A*_1_, again confirming phase alignment.

### anti-phase vibrations

Two waveforms are *anti-phase* if their envelopes exhibit a phase difference of *π*, such that when one increases, the other decreases. This occurs when the envelopes are related through a negatively proportional transformation under a strictly increasing odd function. In this case, the perceived direction of motion alternates across each half-cycle,creating a bouncing or bidirectional trajectory. Perceptually, such out-of-phase vibration patterns may give rise to *bistable motion perception*, wherein the ambiguous temporal dynamics support two competing interpretations, each of which corresponding to motion in opposite directions, that may alternate spontaneously over time. Examples of out-of-phase configurations are shown in Fig. 1D and E.

### Circular motion of a vibrating source

Consider a point source moving along a circular trajectory with radius *r*_0_ and constant tangential velocity *v* (Fig 1C). The frequency of motion is

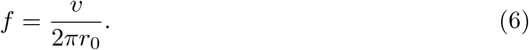

Let *T* be a touch point located at polar coordinates (*r, φ*) relative to the centre of the circular path *O*. The received vibration at time *t* is given by:

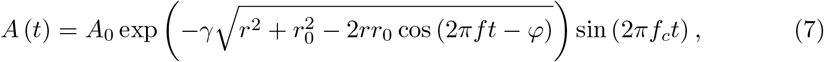

where *A*_0_ is the source amplitude, *f*_*c*_ is the carrier frequency, and *γ* is the attenuation coefficient of the medium. The *envelope* of the received vibration is the time-varying function:

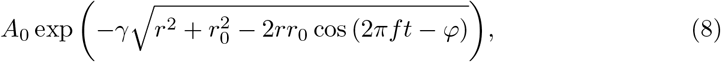

which oscillates at frequency *f*, between a minimum of *A*_0_ exp (−*γ* (*r* + *r*_0_)) and a maximum of *A*_0_ exp (−*γ* |*r* − *r*_0_|).

### Extension to 3D

In three dimensions, the equation for the received signal becomes:

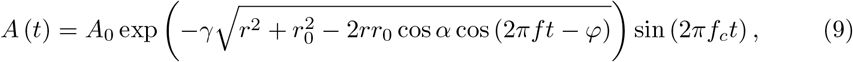

where *α* is the elevation of the touch point *T* relative to the trajectory plane denoted by *P*, and 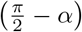 is the inclination angle in spherical coordinates (Fig 1C).

### Phase Differences from Geometry

Let *P* denote the plane of circular trajectory, centred at *O*. Consider two touch points, *T*_1_ and *T*_2_, with projections 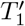 and 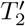 onto plane *P*. Without loss of generality, let the coordinates of the two touch points be (*θ*_1_, *φ, r*_1_) and (*θ*_2_, 2*π − φ, r*_2_), respectively.According to Eq. 9, the received vibrations at the two touch points are:

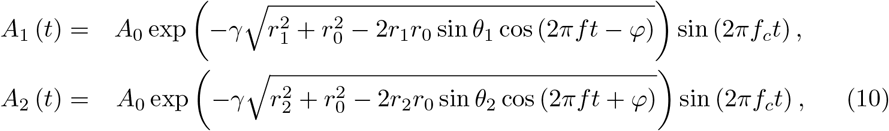

Thus, the envelope phase difference between the two points is Δ*φ* = 2*φ*.

Assume that the projected points and the centre of the circular path *O* lie on a straight line (Fig 1D). Then, if *O* lies **between** 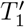 and 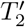 (i.e., 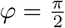) the envelopes of the vibrations are **anti-phase** (Fig. 1D and E). Conversely, if *O* lies **outside** the segment connecting 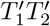– i.e., *φ* = *π*,– the envelopes are **in phase**.

For simplicity, hereafter, we focus on the 2D symmetric case where the centre of the circular trajectory lies on the perpendicular bisector of the segment connecting the two touch points *T*_1_ and *T*_2_, such that *r*_1_ = *r*_2_. In this configuration, the two touch points are equidistant from the centre, resulting in envelopes of vibration with equal amplitude range. This condition facilitates visualisation and analysis of in-phase and out-of-phase conditions in a two-dimensional geometry.

## Experimental procedure

Three psychophysical experiments were conducted to investigate vibrotactile motion direction discrimination using a common two-alternative forced-choice (2-AFC) discrete trial paradigm. In this experiment, participants reported the perceived direction of vibro-tactile motion (left vs. right) generated by two amplitude-modulated vibrations delivered simultaneously to the index and middle fingertips of the right hand (see details below). All experimental procedures were approved by the Monash University Human Research Ethics Committee (MUHREC) and conducted in accordance with approved guidelines.

### Participants

A total of 26 participants (12 female, age range: 19–34, 1 left-handed) took part across the three experiments. All were undergraduate or graduate students at Monash University. Each experiment involved distinct participant groups. All participants provided written informed consent prior to the experiment.

### Vibro-tactile stimulation

In all experiments, vibrotactile stimuli were delivered simultaneously to the index and middle fingertips of the right hand using two miniature solenoid transducers (PMT-20N12AL04-04, Tymphany HK Ltd; 4 Ω, 1 W, 20 mm diameter) mounted 5 cm apart on a vibration-isolated pad. Stimuli were generated in MATLAB (MathWorks Inc.) at a sampling rate of either 48 kHz or 192 kHz, and output through a Creative Sound Blaster Audigy Fx 5.1 sound card (model SB1570). The peak-to-peak amplitude of the output waveform was set to 1.98 V. The shape and curvature of the transducer matched the size and contour of adult fingertips [28]. The base (carrier) frequency of the vibrations was *f*_*c*_ = 100*Hz*. Although this frequency is within the audible range, we verified during pilot testing that the stimuli were not audible to participants and could only be perceived via touch. Amplitude modulation (AM) was applied to generate low-frequency envelopes, with each trial containing 3 modulation cycles. The modulation amplitude was set well above the detection threshold, and pilot testing confirmed that even halving the amplitude had negligible effects on performance in the motion discrimination task.

In all experiments, sinusoidal envelopes were used due to their mathematical and physical properties. Sinusoids are fundamental in Fourier decomposition and are the only waveforms that preserve their shape under summation with others of the same frequency. Sinusoidal modulation mimics natural oscillatory signals (e.g., wind, light, and sound waves) and implies motion with varying velocity, similar to pendular or spring-mass dynamics.

For a given phase difference Δ*φ*, the two sinusoidally modulated vibrations were defined as:

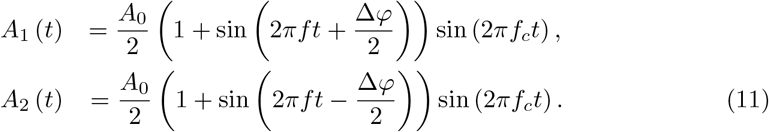

To avoid any response bias or cues about motion direction arising from differences in initial envelope amplitude, vibration onset was set to one of the two isoamplitude points where the envelopes were identical. For non-zero Δ*φ*, these occur at 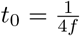 and 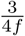 with the corresponding envelope amplitudes of 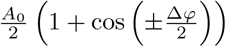 and 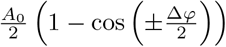, respectively (Fig 2C). Since cosine is an even function, the envelope magnitudes are identical for +Δ*φ/*2 and −Δ*φ/*2. At each of these onset points, the envelopes have opposite slopes – one rising and the other falling – corresponding to opposite directions of motion along the circular trajectory. On each trial, one of these two onset points was selected at random with equal probability, ensuring that initial envelope phase provided no reliable cue about motion direction.

**Fig 2.**
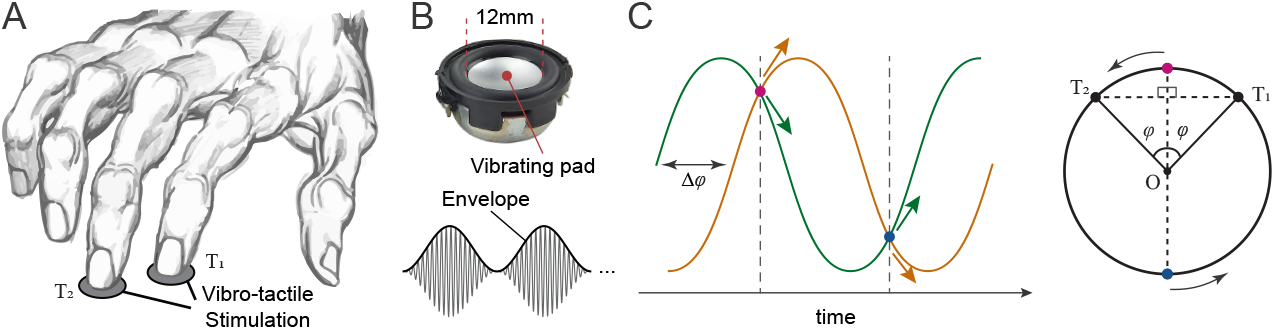
Motion direction discrimination task. **A**, The index and middle fingers of the right hand were stimulated using a pair of solenoid transducers (upper panel, **B**), which delivered amplitude-modulated vibrations (lower panel, **B**). **C**, On each trial, the envelopes of the two vibrations had a phase difference Δ*φ*. The vibrations began at one of two points where their envelope amplitudes were equal (indicated by dashed lines).

In Experiment 1, we additionally included stimuli with exponentially decaying envelopes to simulate more realistic, physically plausible patterns of vibration propagation. The envelopes were derived from Eq. 10, using fixed parameters *r*_0_ = *r*_1_ = *r*_2_ = 1, and 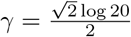:

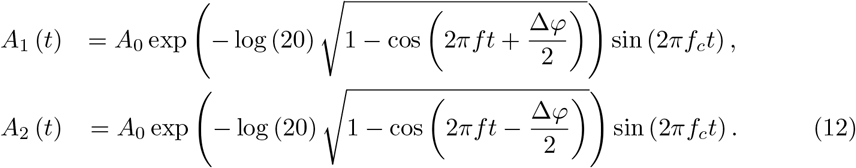

These stimuli were also initiated at one of the two isoamplitude points selected randomly on each trial, consistent with the sinusoidal condition.

## Motion direction discrimination task

Participants performed a discrete-trial two alternative forced-choice (2-AFC) task to judge the perceived direction of vibrotactile motion. On each trial, two amplitude-modulated vibrations were delivered simultaneously to the index and middle fingertips of the right hand. Participants were instructed to gently rest their fingertips on the transducers without applying force (Fig. 2). The two transducers were spaced 5 cm apart on a vibration-isolated pad, arranged such that vibrations from one transducer were not perceptible at the other. Participants rested their arm on the chair armrest with their wrist comfortably supported on a padded surface aligned with the stimulation platform. They were instructed to maintain a stable hand posture throughout each session. All participants reported clear perception of the envelope modulation, and pilot testing confirmed that the stimulus amplitude was well above the detection threshold.

The task was self-paced, with all responses made via keyboard. On each trial, a pair of vibrations with a specific envelope phase difference was presented for three cycles (e.g., 6 s at an envelope frequency of 0.5 Hz). Participants reported the perceived motion direction (leftward or rightward) by pressing the corresponding arrow key with their left hand. There was no time limit for responses, and participants could respond at any moment during or after stimulation.

The specific phase differences and envelope modulation frequencies varied across the three experiments. In Experiment 1, we compared sinusoidal and exponential envelopes with phase differences of ±30°to ±150°(30° increments) at a fixed envelope frequency of 0.5 Hz. In Experiment 2, sinusoidal envelopes were used with phase differences ranging from -180° to 180° in 30°increments, tested at envelope frequencies of 0.5, 1, and 1.5 Hz. In Experiment 3, sinusoidal envelopes were tested at phase differences of 0°, ±30°, ±60°, ±90°, and 180° at a fixed frequency of 0.5 Hz, with participants additionally providing confidence ratings after each response by pressing a number key from “1” (no confidence) to “5” (absolute certainty). These ratings were linearly scaled to a 0–100% confidence range. Experimental conditions were presented in pseudorandom order across trials, with approximately 30 repetitions per condition.

## Psychometric Modelling

To quantify sensitivity to phase differences at each modulation frequency, we modelled perceptual discrimination performance as a function of phase difference Δ*φ* using a nonlinear periodic-sigmoid psychometric function:

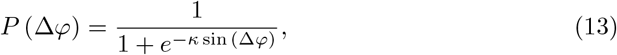

where *P* (Δ*φ*) denotes the predicted proportion of correct responses at a given phase difference Δ*φ*, and *κ* is a sensitivity parameter reflecting the steepness of the psychometric function. Higher *κ* values indicate greater sensitivity to phase differences. This model captures the periodic geometry of the stimulus, predicting chance-level performance (*P* = 0.5) at 0° and 180°, where the vibrations provide no directional cue. The sinusoidal form, combined with its single-parameter structure, avoids overfitting and reflects the hypothetised mechanism of motion perception based on phase-difference readout across spatially separated tactile sensors. The model was fit to group-averaged accuracy data using nonlinear least-squares regression, and the model performance (goodness-of-fit) was assessed using the coefficient of determination (*R*^2^).

## Probabilistic Model of Temporal Reference Detection and Cycle Disambiguation

Consider two AM vibrations wth phase difference Δ*φ*, which corresponds to a temporal lag *d* between their envelopes:

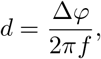

where *f* is the envelope modulation frequency. Let *t*_1_ and *t*_3_ denote the perceived moments of two consecutive salient reference points (e.g., peaks) of vibration 1, and let *t*_2_ denote the corresponding reference point of vibration 2 that occurs between *t*_1_ and *t*_3_. These reference points are extracted from the envelopes of the amplitude-modulated vibrations. Hereafter, we focus on peak features, but the same logic applies to other amplitude landmarks (e.g., troughs or zero-crossings). Due to sensory noise and perceptual limits, each detected peak is assumed to lie within a temporal uncertainty window around the true peak time. For each reference point, I model the perceived time as being uniformly distributed within a window of width *w* centred at the true peak. This uncertainty window depends on the perceptual threshold with which the envelope is extracted. For instance, assuming a sinusoidal envelope in Eq. 11, the time intervals where the envelope deviates from the peak amplitude by less than a threshold value *δ* correspond to durations satisfying |*A*_*i*_(*t*) − *A*_0_| ≤ *δ*, which implies:

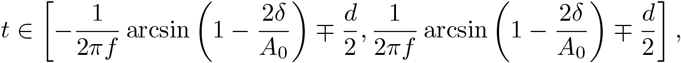

so that the total uncertainty window is:

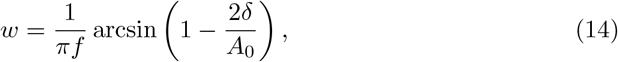

Since the vibrations are periodic and have identical envelope shapes (with vibration 2 being a phase-shifted version of vibration 1), the reference point of vibration 2 is shifted by lag *d*. Similarly, *t*_3_ is one cycle after *t*_1_, i.e., with an offset of *T*, where 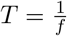 is the envelope modulation period. Without loss of generality, we assume *d >* 0, and align the reference points relative to zero and define their distributions as *t*_1_ ∼ *U* [0, *w*], *t*_2_ ∼ *U* [*d, d* + *w*] and *t*_3_ ∼ *U* [*T, T* + *w*]. Correct perception of motion direction depends on both (1) the reliability of judging the temporal order of salient reference points (e.g., peaks) and (2) disambiguation of within-cycle versus across-cycle intervals. For now, we focus on peak features, but the same logic applies to other amplitude landmarks (e.g., troughs or zero-crossings).

Based on these distributions, the two forms of perceptual inference are required to judge motion direction: first, the **temporal order judgement**, i.e. determining whether the peak of vibration 2 occurs after the peak of vibration 1 (*t*_2_ *> t*_1_). Second, the **inter-peak interval discrimination**, i.e., comparing whether the interval between *t*_1_ and *t*_2_ is shorter than the interval between *t*_2_ and *t*_3_, i.e., testing whether *t*_2_ − *t*_1_ *< t*_3_ − *t*_2_.

The sections that follow formalise these probabilities and derive an analytical expression for the overall probability of a correct motion direction judgement.

### Temporal order judgement

The first source of error arises from uncertainty in judging the temporal order of peaks. If the temporal lag *d*is smaller than the uncertainty window *w*, the perceived ordering of *t*_1_ and *t*_2_ may be incorrect. The probability that the peak of envelope 1 is perceived before that of envelope 2 (i.e., a correct temporal order judgement) is given by the integral of the joint distribution of *t*_1_ and *t*_2_ over the area *t*_2_ − *t*_1_ *>* 0:

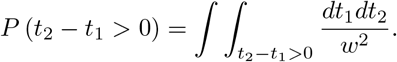

As *t*_1_ and *t*_2_ are mutually independent with uniform distributions, *t*_2_ − *t*_1_ is distributed triangularly over the interval [*d* − *w, d* + *w*], yielding:

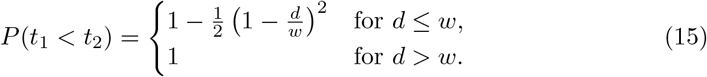

The case*d > w* guarantees correct order due to non-overlapping supports.

### Across-cycle ambiguity and inter-peak interval discrimination

As *d* increases and the uncertainty window extends into the next modulation cycle, another form of error emerges. This is when the uncertainty around *t*_2_ extends beyond the halfway point of the modulation period *T*, the perceived peak of envelope 2 may fall closer in time to the *next* peak of envelope 1 (denoted *t*_3_ ∈ [*T, T* + *w*]) rather than the original one at *t*_1_ ∈ [0, *w*]. This may result in an incorrect interval comparison, i.e.,*t*_2_ − *t*_1_ *> t*_3_ − *t*_2_, thus misjudging the motion direction. Based on the mutually independent uniform distributions of *t*_1_, *t*_2_ and *t*_3_, the probability density function of *V* = *t*_3_ − 2*t*_2_ + *t*_1_ is a piece-wise quadratic function of the form:

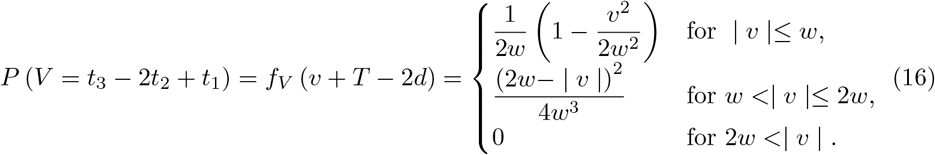

The probability of avoiding this across-cycle confusion is:

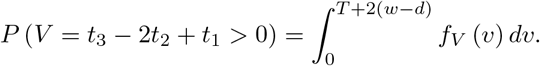

The probability is below 1 when *d* + *w < T/*2.

### Joint probability of correct motion perception

Correct motion perception requires both correct temporal order identification of peaks, and correct across-cycle inter-peak interval discrimination. These two conditions are not independent, and their joint probability must be calculated conditionally. Let Δ = *t*_2_ − *t*_1_. The joint probability of a correct response is:

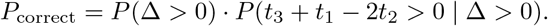

Using the law of total probability over the distribution of Δ, the second term can be rewritten as:

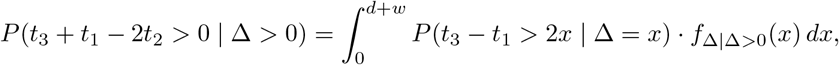

where *f*_Δ|Δ*>*0_(*x*) is the conditional probability density function of Δ = *t*_2_−*t*_1_ given Δ *>* 0, defined as:

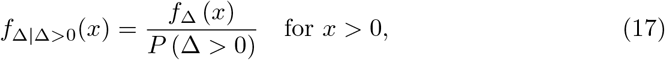

with *f*_Δ_ denoting the triangular probability density function of Δ ∈ [*d*− *w, d* + *w*], and the normalisation constant *P* (Δ *>* 0) as derived in Eq. 15. Thus, the joint probability becomes:

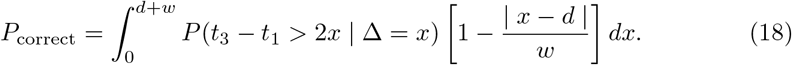

The conditioned probability *P* (*t*_3_ − *t*_1_ *>* 2*x* | Δ = *x*) can be expressed as:

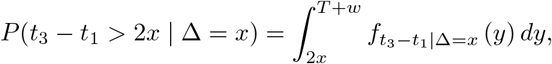

where 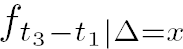 is the distribution of the difference between *t*_3_ ∼ *U* [*T, T* + *w*] and *t*_1_, given Δ = *x*.

The conditional distribution *t*_1_ | (Δ = *x*) ∼ *U* [*a*(*x*), *b*(*x*)], where:

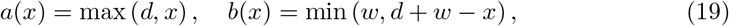

so that the support length is *w*(*x*) = *b*(*x*) − *a*(*x*) = *w* − |*d* − *x*|, at most *w*. This leads to a trapezoidal distribution for *t*_3_ − *t*_1_ | Δ = *x*, from which we derive the cumulative probability:

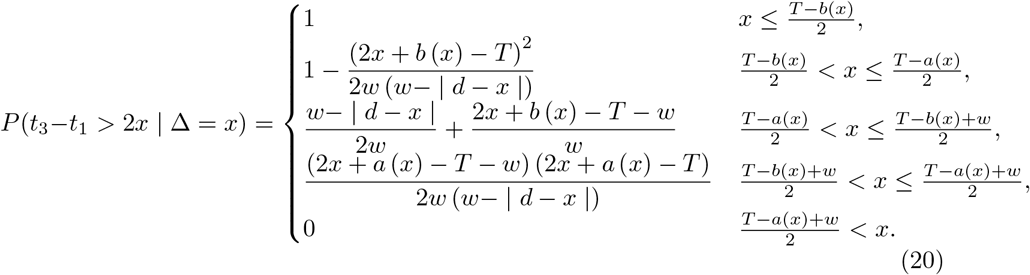

The total probability of a correct decision is obtained by substituting this into Eq. 18. The full expression combines a triangular distribution for Δ, a trapezoidal distribution for *t*_3_ − *t*_1_, and a conditional integration over all valid Δ ∈ [0, *d* + *w*].Though based on simple assumptions, this model predicts a nonlinear psychometric curve that captures key features observed in our data, including asymmetries in performance (e.g., better performance at 30° than at 150° phase lags in Experiment 2).

## Results and Discussion

### Experiment 1: Direction discrimination using sinusoidal vs. exponential envelopes

To examine whether tactile motion perception can arise from simple envelope phase differences alone, I first tested whether participants could discriminate the direction of motion from two simultaneous vibrations with either sinusoidal or naturalistic – i.e., exponential – amplitude modulated (AM) envelopes. Both envelope types simulated a virtual motion trajectory via systematic phase differences across two fingertips.Participants performed a 2-AFC motion direction discrimination task at an envelope frequency of 0.5 Hz. On average across subjects (n = 8), direction discrimination accuracy was 85.5% ±4.2% SEM for exponential envelopes, and 80.2% ±5.1% SEM for sinusoidal envelopes (Fig. 3). While exponential envelopes yielded slightly higher accuracy by 5.3% ±1.5% SEM – possibly due to their closer resemblance to naturalistic wave propagation, – participants still showed robust performance with sinusoidal envelopes. This demonstrates that the tactile system extracts directional information purely from sinusoidal phase offsets, despite their more abstract physical basis.

**Fig 3.**
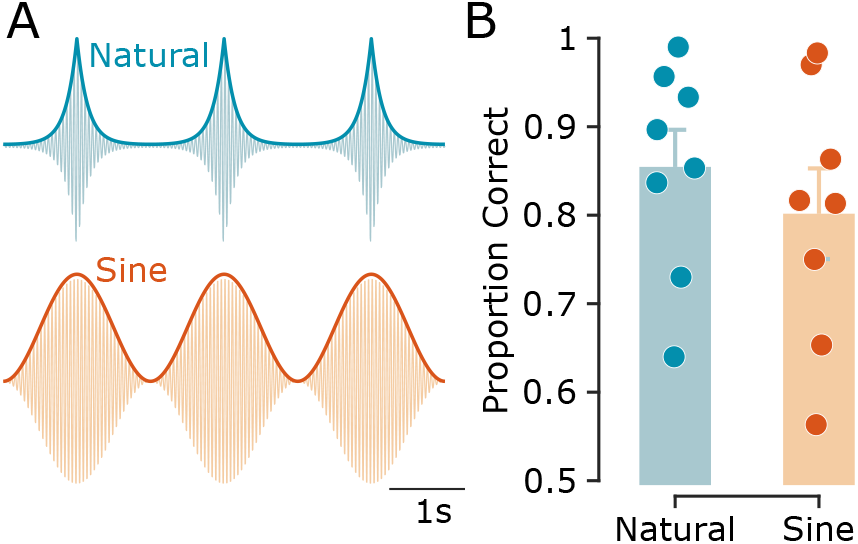
Experiment 1: Naturalistic vs. sinusoidal vibration envelopes. **A**, Schematic representation of naturalistic (exponential) and sinusoidal vibrations, along with their envelopes (thick curves). For illustration purposes, a 20 Hz carrier frequency is shown; the actual carrier frequency used in the experiments was 100 Hz. **B**, Motion direction discrimination accuracy, shown as the proportion of correct trials for exponential and sinusoidal vibrations. Bars represent the average across subjects, with error bars indicating the standard error of the mean (SEM). Data points represent individual participants (*n* = 8).

### Experiment 2: Upper frequency limit for tactile motion discrimination lies below 1.5 Hz

To quantify discrimination performance at each envelope frequency, we fit a sigmoid psychometric model with a sensitivity parameter *κ* to the proportion of correct responses as a function of phase differences (see Methods). At 0.5 Hz, the model fit was robust (coefficient of determination *R*^2^ = 0.56) with a relatively high sensitivity parameter *κ* = 1.29, predicting a maximum accuracy of 78.4%. At 1 Hz, performance declined moderately (*R*^2^ = 0.82, *κ* = 0.78), with a predicted maximum accuracy of 68.5% correct responses. At 1.5 Hz, performance approached chance level (*R*^2^ = 0.36, *κ* = 0.17) with a predicted maximum of just 54.4% correct (Fig. 4), indicating that the upper temporal limit for perceiving direction of tactile motion lies below 1.5 Hz.

**Fig 4.**
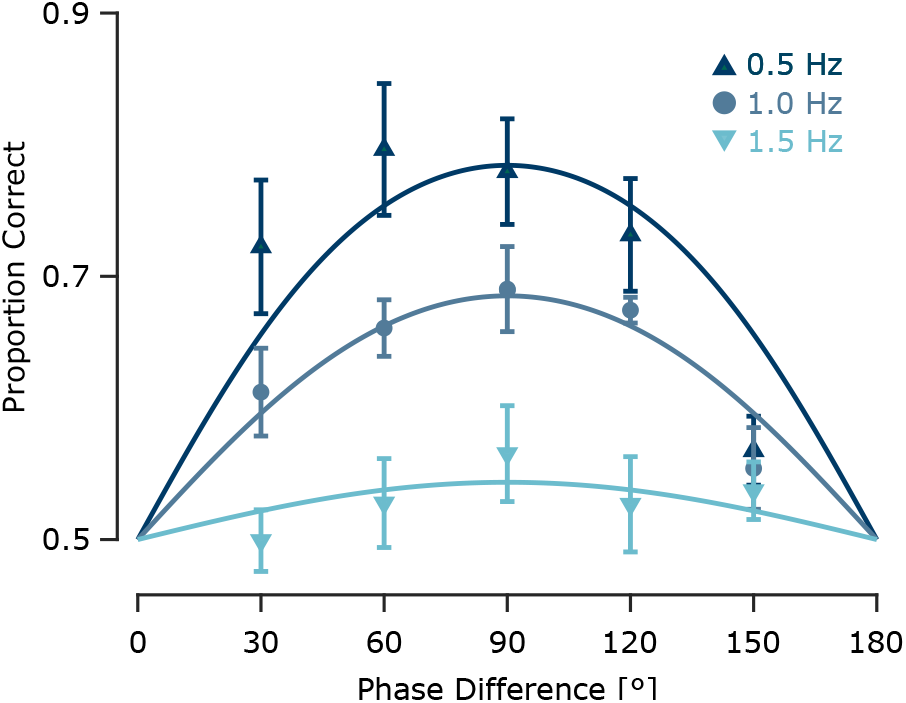
Experiment 2: Effect of envelope frequency on tactile motion perception. Motion direction discrimination performance as a function of phase difference, shown separately for each envelope frequency (indicated by colour). Data points represent across-subject averages (*n* = 6), with error bars indicating the standard error of the mean (SEM). Curves represent psychometric fits for each frequency condition.

These findings contrast with those of Kuroki and colleagues (2016), who examined human sensitivity to AM vibrotactile stimuli up to 20 Hz in a synchronisation-asynchronisation detection task [25]. They reported lower detection threshold frequencies, indicating that participants could reliably detect synchrony or asynchrony in AM signals at nearly ten times higher frequencies than those supporting motion direction discrimination in our study. Moreover, they reported an inverse relationship between modulation frequency and detection threshold, with higher frequencies yielding better synchrony detection. This discrepancy underscores a critical distinction: while humans are capable of detecting synchrony in high-frequency AM signals, perceiving directional motion from inter-finger phase differences relies on much lower envelope frequencies. These differences point to potentially distinct neural mechanisms supporting temporal coincidence detection versus motion perception in the tactile domain.

### Experiment 3: Cognitive and metacognitive signatures of tactile motion perception

Building on Experiments 1 and 2, which established that tactile motion perception depends on the phase difference between fingertip vibrations, Experiment 3 introduced confidence ratings and examined behavioural signatures of perceptual ambiguity. By focusing on phase conditions with minimal directional information (e.g., 0° and 180°), Experiment 3 aimed to characterise how motion uncertainty is reflected in decision confidence, reaction times, and potential choice biases.

#### Ambiguity at 0° and 180° revealed by choice distribution

To assess whether phase differences between fingertip vibrations generate a reliable perception of motion direction, I examined participants’ choices across the range of phase offsets. For directional phase differences (e.g., ±30°, ±60°and ±90°) performance accuracy captures the extent to which participants reported motion direction consistent with the sign of the phase difference (see Fig. 5A). However, at 0° and 180°, the vibrations were either perfectly in-phase or anti-phase across the two fingertips, resulting in symmetric temporal envelopes with no consistent directional cue. As such, for these two conditions, “correct” or “incorrect” responses are undefined for these conditions. Thus, I instead analysed these conditions in terms of choice likelihood – specifically, the proportion of “leftward” responses (Fig. 5B).

**Fig 5.**
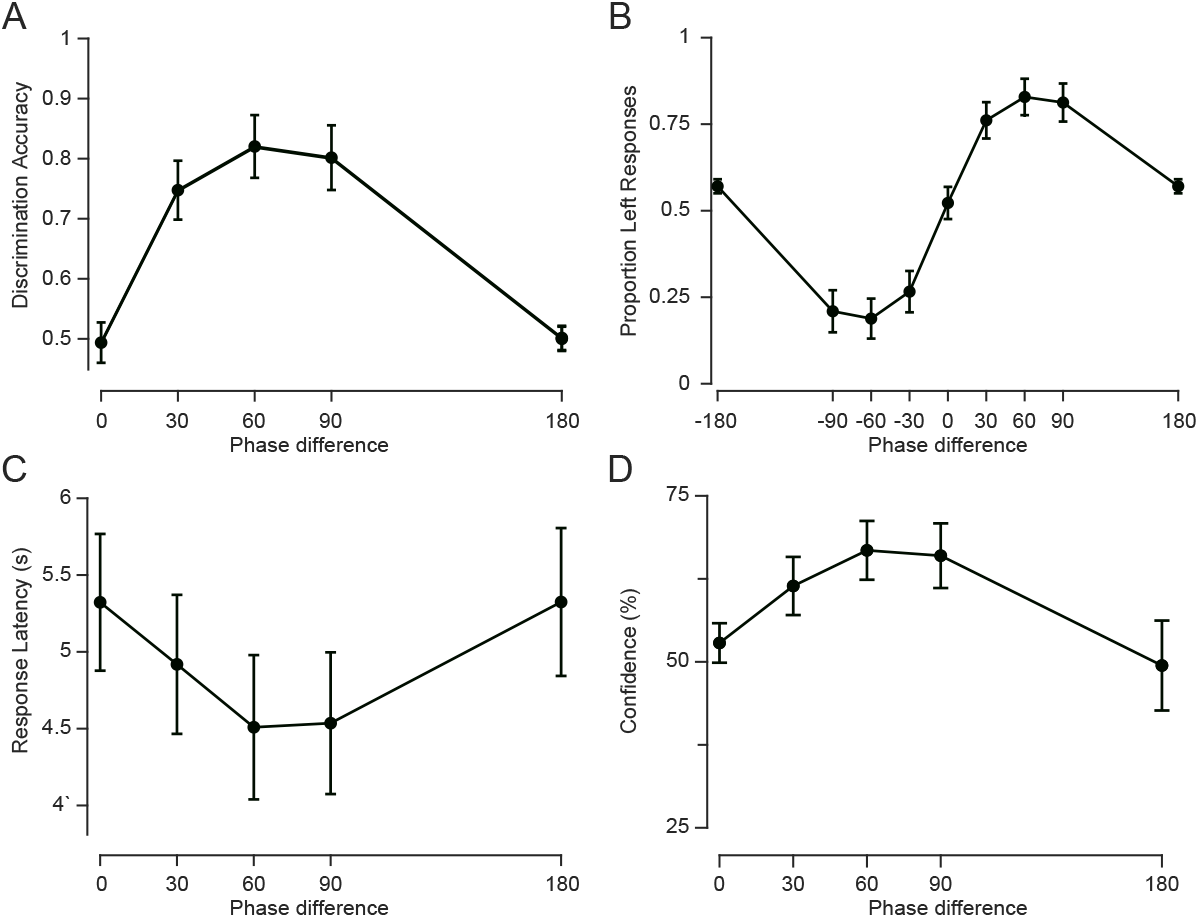
Experiment 3: Cognitive and metacognitive measures of tactile motion. **A**, Discrimination accuracy, measured as the proportion of correct responses, averaged across subjects (*n* = 12). For 0° and 180° phase differences, responses were pseudo-randomly labelled as correct or incorrect. **B**, Psychometric curves (choice likelihood) showing the proportion of “left” responses as a function of phase difference, averaged across subjects. **C**, Median reaction time (interval between stimulus onset and response), averaged across subjects, plotted as a function of phase difference. **D**, Average confidence ratings vs. phase differences. All error bars represent the standard error of the mean (SEM).

At 0°, participants selected “left” on 52.2% of trials (SEM = 4.7%), not significantly different from chance (t(11) = 0.48, p = 0.64), consistent with perceptual ambiguity. At 180°, however, participants showed a subtle but reliable leftward bias (mean = 56.8%, SEM = 2.1%), which was significantly above chance (t(11) = 3.24, p = 0.008). This bias suggests that at 180°, even in the absence of reliable directional cues, early envelope asymmetries or internal decision biases may influence motion judgements.

#### Slower responses reflect ambiguity in motion signal

Response time (RT) provides a behavioural index of the strength of sensory evidence. Here, I analysed how RT changed as a function of phase difference to assess how tactile motion signals support directional judgements under varying degrees of ambiguity. As shown in Fig.5C, RTs followed a U-shaped pattern: responses were slower for the ambiguous conditions (0° and 180°) and faster for intermediate phase differences. To statistically assess this pattern, a linear mixed-effects model (with random intercepts per subject) was fitted to trial-level RTs. Trials with excessively long RTs (*>*15s) were excluded, and the remaining RTs were z-scored within subject to account for individual baseline differences. The model included phase difference and its square as fixed effects. It revealed a statistically significant quadratic effect (*β* = −0.0016, *p<*1e-11),, consistent with an inverted-U pattern of RTs across phase differences, indicating that participants responded more slowly at both 0° and 180° phase differences, and more quickly for intermediate values (30°–90°), as shown in Fig. 5C. This trend is consistent with the interpretation that extreme phase differences (0°, 180°) produce ambiguous or conflicting motion cues, requiring longer processing times. Indeed, the average RTs at 0° and 180° were nearly identical (5.32 ± 0.44 s and 5.32 ± 0.48 s, respectively), and both were higher than for other phase differences.

#### Confidence ratings track motion signal strength

Confidence ratings reflect participants’ subjective assessment of perceptual certainty, providing a metacognitive index of how strongly they perceived the motion signals on each trial. A linear mixed-effects model with subject-wise random intercepts revealed a robust quadratic relationship between confidence and phase difference. Confidence was lowest at extreme phase difference values (0° and 180°) and highest at intermediate phase differences (30°–90°), mirroring the pattern observed in reaction times (Fig. **??**D), and consistent with weaker or more ambiguous motion signals at the extremes. This trend was reflected in a significant negative quadratic term (*β* = −0.0021, *p <* 1*e* −44) and a significant positive linear term (*β* = 0.36, *p <* 1*e* −35). These results indicate that participants were more confident when phase differences provided stronger directional cues, and less confident when the motion signal was more ambiguous.

In particular, average confidence at 180° was 49.4% ±6.8% (mean ± SEM across participants), lower than all other conditions, including0° (52.8% ±3.0%), supporting the interpretation that anti-phase stimulation elicits especially uncertain percepts.

### Phase differences, not amplitude differences, drive tactile motion perception

The behavioural ambiguity observed at 180° phase difference – reflected in a subtle directional bias, low confidence, and slow responses – raises a critical question: What stimulus features underlie tactile motion perception? Two plausible mechanisms are: (i) motion perception based on the *phase difference* between two AM signals (i.e., relative *temporal shifts* in their envelopes), and (ii) perception based on *moment-by-moment amplitude differences* between the signals.

The present stimulus design enables these alternatives to be dissociated. While ±180° phase differences produce the largest instantaneous amplitude differences between fingertips, they contain no consistent directional information, as the +180° and -180° stimuli are physically identical and indistinguishable. If motion perception were driven by amplitude differences alone, one would expect robust and consistent directional judgements under these conditions – contrary to the observed choice likelihood patterns.

Moreover, an amplitude-based account might predict high confidence on individual trial bases (despite random direction across trials), assuming a salient motion signal. Yet, confidence ratings at 180° were the lowest across all phase differences, mirroring the slower responses typically associated with perceptual uncertainty. Together, these findings support a mechanism in which *phase differences* between signals, not momentary amplitude (or “enegry”) differences, drive tactile motion perception.

### Potential underlying neural computations

As in vision, tactile motion perception may rely on multiple neural computations [5, 29]. Here, I outline two candidate mechanisms that could support the perception of motion based on phase differences in vibrotactile signals. These mechanisms differ in whether they rely on the measures of similarity of temporal patterns or on the relative timing of specific features (e.g., peaks or troughs) in the tactile signals. Below, we briefly discuss each and assess their neural plausibility.

#### Temporal cross-correlation mechanisms

A plausible computational mechanism underlying tactile motion perception is based on temporal cross-correlation of the continuous tactile sensory inputs received at the two fingers. In this scenario, the brain compares the envelopes of each vibration over a certain temporal window to estimate their relative lag (phase difference) similar to Reichardt detectors [1, 30]. The inferred phase Neural mechanisms for such temporal cross-correlation have been widely studied in other sensory systems. For example, in the auditory system, interaural time differences are computed via temporally sensitive circuits in the medial superior olive, involving coincidence detection mechanisms [31]. In the electrosensory system of weakly electric fish, neurons perform delay-sensitive comparisons between signals from different electroreceptors to extract motion or phase differences of preys [32]. While mammalian tactile system may not contain dedicated delay lines, some neurons in somatosensory cortex (particularly S1 and S2) exhibit phase-locked responses to frequency modulations [33, 34], carrying information about the temporal patterns of sensory inputs. Additionally, cross-digit integration occurs at multiple levels, including primary and secondary somatosensory cortices, where receptive fields often span multiple fingers [35, 36]. Such distributed, temporally sensitive representations could support correlation-based decoding of phase relationships. The observed sensitivity to small phase differences (e.g., 30°) in the present study is consistent with this type of integration. Thus, a biologically plausible hypothesis is that populations of neurons in somatosensory cortex, or possibly parietal areas, integrate envelope information and compare their temporal alignment. Population-level decoding of such temporal relationships could underlie the perceptual sensitivity to direction based on phase difference, as observed in our experiments. Whether these computations occur via explicit cross-correlation at the neural level, or are approximated by population-level pooling across temporal patterns, remains to be clarified.

Importantly, these computations are not limited to biological intuition but are also grounded in formal estimation theory. Under assumptions of linearity and Gaussian noise, cross-correlation, least-squares, and maximum likelihood methods yield equivalent estimates for time delay between signals [37]. These mechanisms are sensitive to the overall similarity and alignment of time-varying signals, rather than to discrete features such as peaks or zero-crossings. As such, they can operate continuously and flexibly, and do not depend on precise extraction of singular time points, potentially making them robust to noise.

#### A probabilistic model from envelope landmarks

A second mechanism is that the tactile system detects specific temporal landmarks in the envelope of each vibration – such as peaks, troughs, or other salient features – and infers motion direction based on the temporal order or timing of these events relative to each other. This process is inherently susceptible to sensory variability (noise) and perceptual uncertainty, especially when the modulated envelope changes gradually or when features are close in time.

To formalise this temporal uncertainty, I propose a simple threshold-based model in which a temporal reference point (e.g., a peak) is detected when the change in the envelope exceeds a certain slope or amplitude threshold. Changes below this threshold are not perceived as distinct events. For instance, under this assumption, any portion of the envelope around the true peak whose amplitude lies within the threshold margin is perceptually indistinguishable from the true peak. This introduces variability in the perceived timing of features or leads to missed detections, particularly when the modulation depth is shallow or the envelope is slowly varying.

Such detection uncertainty can lead to errors in temporal order judgements. For instance, two peaks occurring closely in time might be perceived in the wrong order, or the tactile system might match a peak from one vibration to the wrong cycle of the other, especially under large phase differences. These errors impair the brain’s ability to reliably infer motion direction. Importantly, this minimal model – based on a fixed amplitude detection threshold without any complex decoding and uniform temporal variability – produces non-trivial psychometric predictions. As illustrated in Fig. 6, the model generates an asymmetric curve of predicted proportion correct as a function of phase difference: performance increases steeper near small positive phase offsets (e.g., 30°), but declines more gradually beyond 90°, reflecting cross-cycle misalignments and detection failures (e.g., for 150°). This asymmetry is evident in our experimental data, particularly in Experiment 2, where performance at 30° phase difference is higher than at 150°, despite the physical symmetry of the stimuli.

**Fig 6.**
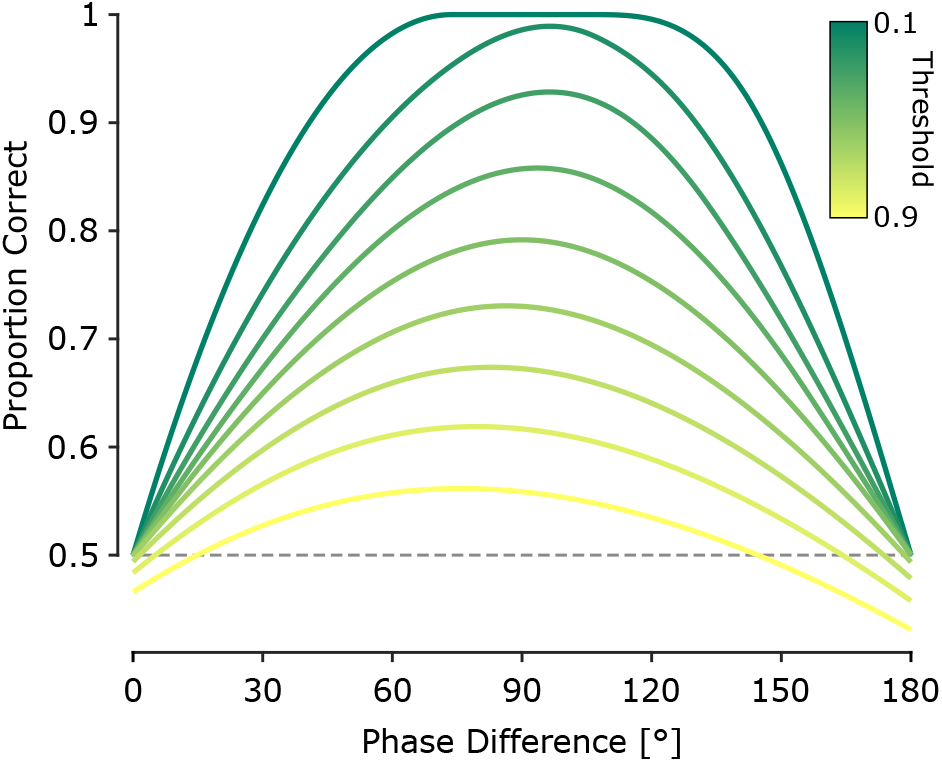
Predicted direction discrimination performance from the probabilistic feature-based model. Model-predicted proportion of correct motion direction discrimination as a function of phase difference. Each trace corresponds to a different amplitude detection threshold (indicated by colour), expressed as a proportion of the peak envelope amplitude (from 0.1 to 0.9 in increments of 0.1).

While this model was implemented based on peak detection, the same logic applies to other types of envelope features, including troughs or points of inflection. The key principle is that temporal reference points are perceived only if they exceed a salience threshold, and that perceptual errors emerge from variability in the timing or detectability of these points. This model captures the dual sources of perceptual error: (1) local ambiguity in temporal order when the temporal reference points are too close, and (2) misattribution across cycles when phase differences approach 180°.

Critically, this model explains why perceptual performance deteriorates at large phase differences despite increased amplitude contrast: the temporal lag between reference points (e.g. peaks) is closer to *T/*2, increasing the probability that reference points from one vibration are misattributed to a different cycle of the other. These findings suggest that under threshold-limited temporal resolution, tactile motion perception involves a delicate balance between fine temporal discrimination and the global temporal structure of the stimulus.

Together, these results suggest that tactile motion perception across fingers could be shaped by both temporally global (e.g., cross-correlation) and local (event-based) temporal processing mechanisms, each with distinct neural constraints and noise profiles.

## Conclusion

This study investigated how the tactile system extracts spatial information about object motion from temporally structured vibrations delivered to two fingertips. Across three experiments, I delivered pairs of amplitude-modulated vibrations – each comprising a 100 Hz carrier modulated by a low-frequency sinusoidal envelope – to simulate *continuous* tactile motion. By systematically varying the phase difference between the two envelopes, I quantified how inter-fingertip phase offsets influence perceived motion direction, response latency, and confidence.

Our findings confirmed that the direction of perceived motion is determined by the phase difference between the two vibrations and not by their absolute frequency or amplitude. Experiment 1 showed that sinusoidal envelope vibrations reliably elicited robust directional motion percepts, comparable to those evoked by natural patterns (e.g., exponential decay). Notably, [12] found that gradually ramped vibrotactile stimuli produced stronger and smoother motion percepts than abrupt onsets, consistent with the use of continuous amplitude modulated vibrations in the present study to simulate naturalistic motion cues. Experiment 2 established that the upper frequency limit for reliable tactile motion discrimination lies below 1.5 Hz, nearly ten fold higher frequency threshold than those reported in earlier studies using similar stimuli [25]. Experiment 3 revealed systematic changes in confidence and reaction time with phase difference, with ambiguous conditions (0° and 180°) producing slower responses and lower confidence ratings. Importantly, although the 180° condition, despite producing the largest moment-by-moment amplitude differences between fingers, did not yield a consistent percept of direction, suggesting that motion perception depends on phase differences, not amplitude disparity.

Together, these results provide new insight into the computational basis of tactile motion perception. They support a mechanism in which tactile motion perception arises from the relative *phase differences* between of temporally structured signals across skin locations, rather than from instantaneous amplitude differences or ‘energy shifts. Unlike prior studies of tactile synchrony detection, the present paradigm required spatial trajectory inference across inputs, revealing that *phase-based temporal integration*, rather than amplitude contrast, underpins tactile motion perception. While Kuroki et al. (2016) demonstrated that humans can detect temporal asynchrony in AM tactile stimuli at higher modulation frequencies (up to 20 Hz, [25]) indicative of sensitivity to temporal structure, their task probed asynchrony detection, not motion inference.Drawing parallels to the visual system, they proposed that tactile perception may rely on both “phase-shift” and “energy-shift” mechanisms, analogous to first- and second-order motion processing in vision.

Notably, Kuroki et. al reported peak detection at a 180° phase difference. Yet in the current study, the same phase difference produced ambiguous motion percepts, reflected in lower confidence, slower responses, and inconsistent choices. This discrepancy likely reflects task-specific neural computations for synchrony detection and motion perception. Synchrony detection may rely on local temporal contrast or energy cues at single skin locations, whereas tactile motion perception requires spatial comparison and temporal integration across fingertips. The present results suggest that phase-based readout, rather than local amplitude difference, is central to tactile motion perception.

This dissociation highlights that motion perception depends on the integration of temporal phase relationships across space and time. As in the visual system, where distinct pathways support multiple forms of motion processing, the tactile system may also engage parallel mechanisms for temporal analysis. Phase-based computations appear specifically tuned for inferring motion trajectories, distinguishing them from those supporting synchrony detection. These findings reveal how the tactile system transforms temporally structured input into spatial motion percepts, and how the brain selectively engages distinct temporal codes based on perceptual goals.

Here, I proposed two complementary models of tactile motion perception; one based on global cross-correlation of vibration envelopes, and another relying on local temporal comparisons between salient features such as envelope peaks. While the cross-correlation model captures overall waveform similarity, the feature-based model formalises direction perception as a probabilistic judgement derived from uncertain detection of temporal landmarks within amplitude-defined windows. Notably, both models are applicable to conventional apparent motion paradigms, where discrete or pulsed stimuli with staggered onsets simulate movement. Although the inter-peak intervals in our stimuli (e.g., 167 ms for 30° and 500 ms or 90° at 0.5 Hz) exceed classical tactile temporal order judgement thresholds [38], participants nonetheless exhibited robust directional performance and systematic confidence patterns. Notably, performance at 30° phase lag aligns with previously reported temporal order judgement thresholds (∼100 ms, [38]), despite differences in stimulus type and parameters, suggesting that reliable direction perception can emerge without discrete onsets or overt spatial displacement. While supramodal attentional tracking could, in principle, support such judgements – e.g., by tracking salient events across time and space irrespective of sensory modality, – our model provides a tactile-specific alternative. It attributes direction perception to probabilistic comparisons between uncertain temporal landmarks (e.g., envelope peaks), detected within amplitude-defined integration windows. This framework captures This framework captures the nonlinearity in psychometric curves, including both the reliable direction perception at shorter phase lags and the ambiguity at 180°, without invoking higher-level amodal mechanisms or cross-modal attentional strategies. Instead, it reflects constraints intrinsic to tactile processing, where perceptual uncertainty in temporal feature extraction shapes directional judgements.

Central to the perception of the vibration-induced motion studied here is the brain’s ability to track dynamic changes in the envelopes of tactile signals and extract directional information from their relative timing. This sensory strategy has analogues across species: arachnids, for example, detect prey using complex vibration patterns transmitted through webs or substrates, relying on finely tuned mechanosensory systems that evolved independently from vertebrate touch [24]. In mammalian glabrous skin, Meissner’s and Pacinian corpuscles are specialised for detecting vibration [39–43], with Pacinian corpuscles implicated in encoding vibrotactile pitch in both mice and humans [44–47]. My previous work demonstrated that rodents can discriminate vibrations based on both amplitude and frequency using their whiskers [34, 48, 49].Neurons in primary somatosensory cortex integrate these features in a way that supports vibrotactile perception. The present study builds on these principles, showing that temporal features – specifically phase relationships – can be exploited to generate robust perceptions of tactile motion across fingertips. This supports the idea that tactile systems, across species and sensor types, flexibly encode both spectral and temporal properties of mechanical stimuli to extract high-level perceptual content.

A major challenge in studying somatosensation is the ability to deliver tactile stimuli with precise control and reproducibility. In freely moving animals, variations in posture, movement, and skin contact can significantly affect the quality and consistency of tactile stimulation, introducing variability in the sensory input and complicating the interpretation of neural responses. In humans, the elastic properties of the skin can lead to trial-by-trial differences in receptor activation due to subtle changes in pressure, tension, or contact geometry [3]. These inconsistencies may engage different mechanoreceptor subtypes, potentially altering the percept and confounding behavioural measurements. To address these limitations, I developed a vibrotactile stimulation paradigm in which the perception of motion direction is determined not by low-level features of the individual vibrations – such as absolute amplitude or frequency – but by the phase relationship between them. This design enables consistent control over the critical perceptual variable (phase difference), even when some variability in contact conditions is unavoidable. As such, it offers a robust and generalisable framework for investigating tactile motion perception and related decision-making processes.

Finally, this paradigm provides a powerful tool for probing tactile decision-making and perceptual inference under controlled temporal structure, paralleling the role of random-dot motion in visual neuroscience. By dissociating low-level vibration attributes from high-level motion perception, it offers a flexible approach for linking somatosensory encoding with computational models of evidence accumulation and perceptual categorisation, in both human and animal research. By revealing how the brain transforms temporally structured input into coherent motion percepts across the skin, this work contributes to a deeper understanding of somatosensory processing and lays the groundwork for future research in touch-based interfaces, neuroprosthetics, and tactile cognition.

## Acknowledgments

The author thanks Mohammad Razmjoo and Erfan Rezaei for their assistance with data collection in Experiment 1.

